# Bacterial c-di-GMP plays a key role in the evolution of host-association

**DOI:** 10.1101/2023.03.20.533436

**Authors:** Nancy Obeng, Anna Czerwinski, Daniel Schütz, Jan Michels, Jan Leipert, Florence Bansept, Thekla Schultheiß, Melinda Kemlein, Janina Fuß, Andreas Tholey, Arne Traulsen, Hinrich Schulenburg

## Abstract

Most microbes evolve faster than their hosts and should therefore drive evolution of host-microbe interactions^1–3^. However, relatively little is known about the characteristics that define the adaptive path of microbes to host-association. In this study we have identified microbial traits that mediate adaptation to hosts by experimentally evolving the bacterium *Pseudomonas lurida* with the nematode *Caenorhabditis elegans*. We repeatedly observed the evolution of beneficial host-specialist bacteria with improved persistence in the nematode, achieved by mutations that uniformly upregulate the universal second messenger c-di-GMP. We subsequently upregulated c-di-GMP in different *Pseudomonas* species, consistently causing increased host-association. Comparison of Pseudomonad genomes from various environments revealed that c-di-GMP underlies adaptation to a variety of hosts, from plants to humans, suggesting that it is fundamental for establishing host-association.

## Main text

Host-associated microorganisms have important effects on the physiological functioning and fitness of their plant and animal hosts^2, 4^. These host-microbiota interactions are often studied using a host-centric view, with a focus on microbiota-mediated host functions. This view neglects the important fact that most microbes evolve faster than their hosts, due to their shorter generation times and higher mutation rates, and thus that fitness improvements for the microbes may disproportionately drive the associations^1^. An important step in the evolution of a host-microbe association is emergence of a more specialized interaction that allows free-living bacteria to reliably enter the host, persist and finally be released into the environment to colonize new hosts (Fig. 1a, line 102)^1^. To date, little is known about the traits and molecular processes that determine how bacteria adapt to such an association with the host.

**Fig. 1.**
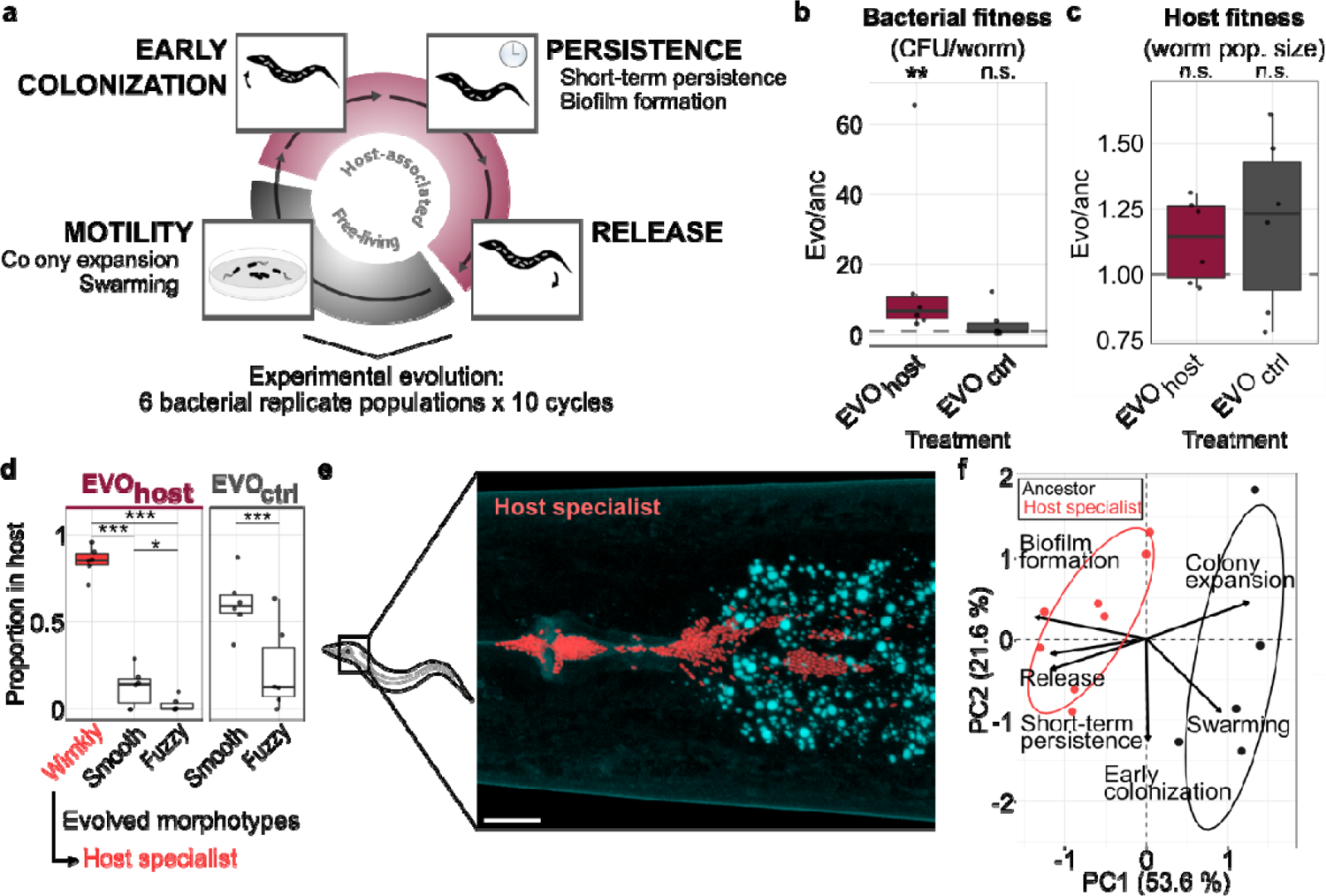
Microbiota bacteria evolve a host-specialist phenotype. **a**, Host-associated microbes transition from a free-living phase to host-association, the latter comprising host entry, persistence, and release. Six *Pseudomonas lurida* populations were passaged ten times across these stages with the host *Caenorhabditis elegans* (EVO_host_) or without the host as a control (EVO_ctrl_). **b**, Host-adapted bacterial populations significantly increased fitness (given as colony forming units CFU/worm) relative to the ancestor (t-tests and Tukey post-hoc comparisons, 6 replicates/treatment). **c**, Evolved bacteria remain beneficial to the host, determined by nematode population growth (t-test, 6 replicates/treatment). **d**, A wrinkly colony morphotype only emerged during host adaptation and dominates within worms (comparison of morphotype abundance within treatments: generalized linear models and Tukey post-hoc test, 6 replicates/treatment). **e**, Evolved host-specialist bacteria (tagged with red fluorescent dTomato) colonize the worm gut (scale bar = 10 µm). **f**, Principal component analysis of key traits of the host-associated life cycle (in **a**) reveals a distinct profile for evolved wrinkly host specialists compared to ancestral bacteria (see Supplementary Table 5 for individual traits measurements). (**b**, **c**, **d**) * = P < 0.05, ** = P < 0.01, *** = P < 0.001, n.s., not significant.

### Evolution of host specialist bacteria

We studied the evolutionary transition from free-living to host-association through controlled experimental evolution, using the bacterium *Pseudomonas lurida* and the nematode host *Caenorhabditis elegans* as a model. This bacterium is occasionally found in the natural microbiota of *C. elegans*^5, 6^. Under laboratory conditions, the presence of *P. lurida* is associated with increased population growth rates of *C. elegans* and it can provide protection against pathogens, yet both host and bacterium can proliferate without each other and thus do not depend upon one another^5, 7, 8^. To select host-adapted bacteria we serially passaged 6 *P. lurida* populations either with or without a host-associated phase (Fig. 1a, line 102; EVO_host_ or EVO_ctrl_, respectively). All populations were inoculated from the same clonal ancestor. After 10 passages through hosts, the bacteria reached on average 5-10 times higher bacterial load in the host than their ancestor, a significant change not observed for the control that evolved without exposure to hosts but otherwise identical conditions (Fig. 1b, line 102, Extended Data Fig. 1). The increased bacterial fitness did not come at a cost to the host, as nematode population growth (used as a proxy for nematode fitness^5^) did not change significantly, but rather increased in the presence of the adapted bacteria (Fig. 1c, line 102).

As a result of passaging, bacterial populations diversified in colony morphology. At the end of our experiment a “wrinkly” morphotype was dominant in all host-associated experimental replicate populations and absent in the controls, whereas “fuzzy” and “smooth” (ancestral) morphotypes were present across treatments (Fig. 1d, line 102, Supplementary Table 1). Despite their significant advantage in hosts, the wrinkly morphotypes declined during growth on agar, while smooth and fuzzy types increased in abundance (Extended Data Fig. 2, line 152, Supplementary Table 2). As the wrinkly types were unique to and reached very high abundance in worm-adapted bacteria, we considered them host specialists. These specialists can be found in clusters within the intestinal tract of the nematode, especially in the anterior and posterior parts (Fig. 1e, line 102, Extended Data Fig. 3). Notably, the evolved wrinkly morphotype is similar to wrinkly *P. fluorescens* that emerge at the air-liquid interface in static microcosms^9^, and to rugose variants of various pathogenic bacteria^10–12^. Our experiments suggest that this morphological change also occurs in beneficial bacteria adapting to host-association. For a further characterization of these adaptations, we focused on 47 clones of the distinct and genetically stable morphotypes (Supplementary Table 3), isolated from the final populations of our evolution experiment.

**Fig. 2.**
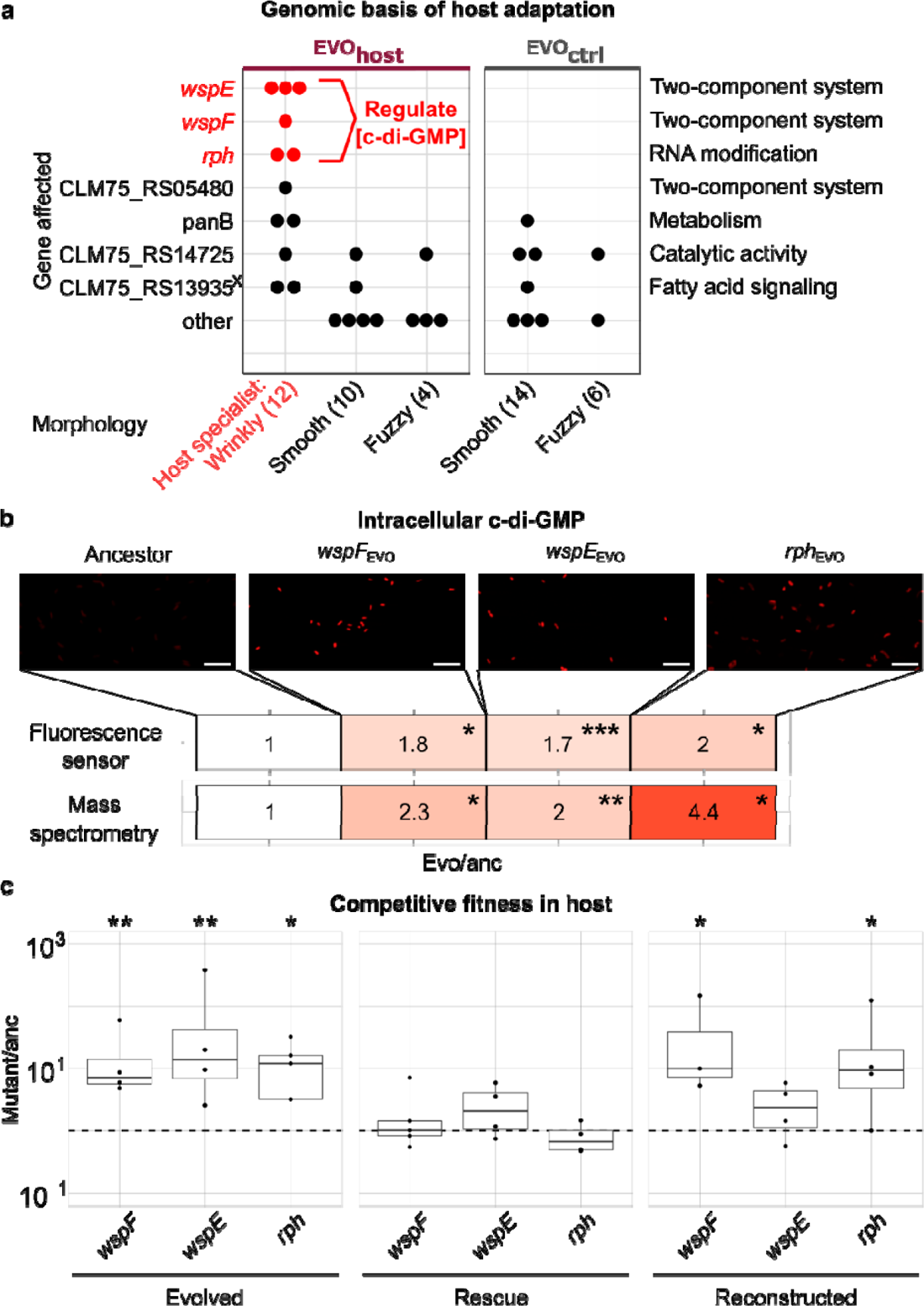
Wrinkly host specialists adapt to *C. elegans* by upregulation of the universal second messenger c-di-GMP and increased intra-host competitiveness. **a**, Overview of genes with non-silent changes in evolved bacterial isolates. Data points represent mutant isolates (total number of sequenced morphotypes in brackets). A cross indicates genes with variants lost in the evolved isolates as compared to the ancestor. **b**, Fluorescence sensor and LC-MS detected higher intracellular c-di-GMP concentrations in evolved wrinkly mutants compared to ancestor (Welch-ANOVA and Games-Howell post-hoc comparisons; scale bar = 10 µm, 5 replicates/treatment). **c**, Competitive fitness (CFU/worm relative to ancestor) of evolved *wspF*, *wspE* and *rph* mutants (left), rescued mutants (middle), and reconstructed mutants in ancestral background (right) during persistence in *C. elegans* (ANOVA and Dunnett post-hoc tests, min. 3 replicates/treatment).; * = P < 0.05, ** = P < 0.01, *** = P < 0.001.

**Fig. 3.**
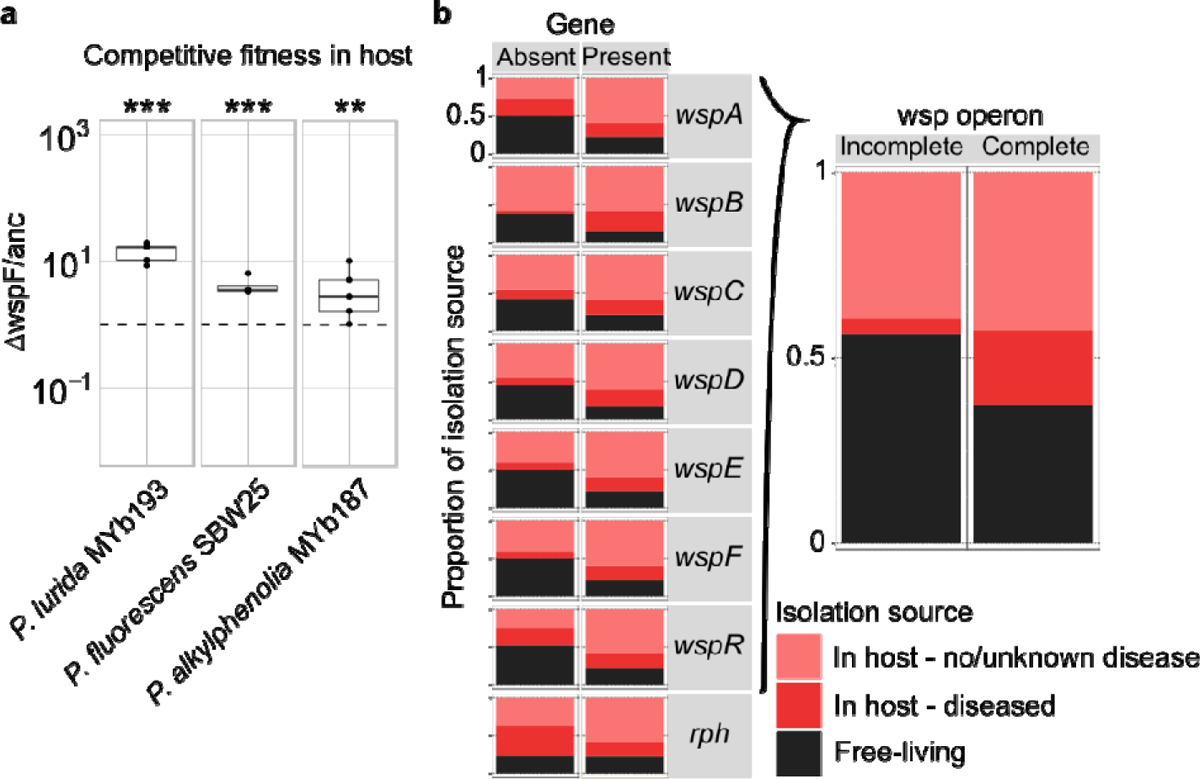
C-di-GMP regulators generally mediate host-association across Pseudomonads. **a**, *WspF* deletion increases intra-host competitive fitness of different *Pseudomonas* species relative to their wildtype (CFU/worm; ANOVA and Dunnett post-hoc tests, min. 3 replicates/ treatment, ** = P < 0.01, *** = P < 0.001). **b**, Presence/absence of *wsp* genes (or operon) and *rph* co-varies with isolation source of sequenced Pseudomonads^2^ goodness of fit tests).

### Host specialist has distinct life style

An analysis of trait changes across the distinct stages of host-association revealed specific adaptations of wrinkly morphotypes to the interaction with *C. elegans*. In detail, we characterized two traits of importance for the free-living stage and four traits for host-association (as listed in Fig. 1a, line 102). We found that the wrinkly isolates were significantly distinct from the ancestral trait profile (Fig. 1f, line 102, Supplementary Table 4). This was mainly due to significant increases in short-term persistence, release from the host, and *in vitro* biofilm formation (Fig. 1f, line 102, Extended Data Fig. 4, Supplementary Table 4, Supplementary Table 5, Supplementary Table 6), all traits that define late phase interactions with the host. The overall pattern of improved host-association was also recovered by analyzing the genetically diverse populations from the end of the evolution experiment, where the host-associated populations similarly increased in persistence and release (Extended Data Fig. 5, Supplementary Table 7, Supplementary Table 8). In detail, biofilm formation can enable persistent contact with the host and increase stress-tolerance^13, 14^, exemplified by many pathogens^15^, thereby improving survival in the nematode’s digestive tract. As a consequence of increased biofilm formation, aggregated cells may be expelled more easily^16^, thereby explaining the observed increase in release. Such shedding also enhances the chance for transmission to other hosts^1^, which restarts the cycle of host-association. Notably, wrinkly isolates did not differ from ancestors in early colonization, yet showed a significant decrease in colony expansion and swarming on plates (Fig. 1f, line 102, Extended Data Fig. 4, Supplementary Table 4, Supplementary Table 5). The latter result is consistent with a decrease in motility described for *E. coli* that evolved to become a mutualist in stinkbugs^17^, but contrasts with findings that sufficient swarming is required for colonization initiation of zebrafish and bobtail squid^18, 19^. These contrasts are likely due to differences in symbiont recruitment between the host systems, defined by either aquatic environments for zebrafish and squid, and terrestrial environments for *C. elegans* and stinkbug, respectively.

Moreover, our observations of increased biofilm formation and reduced motility may indicate an evolved life-history trade-off between the traits defining either host-association and the free-living stage. We conclude that experimental evolution in the presence of the nematode host leads to emergence and spread of a host specialist type. We next asked whether the improved host-association has a common genetic basis.

### c-di-GMP determines host specialization

Whole genome sequencing of the isolated morphotypes and the ancestor revealed several independent mutations in wrinkly host specialists that affect the universal second messenger cyclic diguanylate (c-di-GMP). In particular, a comparison of non-silent genomic variation identified variant genes specific to wrinkly host specialists (Fig. 2a, line 152, Supplementary Table 9). Two of the genes, *wspE* and *wspF*, code for a hybrid sensor histidine kinase and a methylesterase in the wrinkly spreader (*wsp*) operon, respectively^20^. These genes are part of a two-component system that regulates c-di-GMP levels (Extended Data Fig. 6) and wrinkly formation in beta- and gammaproteobacteria, including Pseudomonads^20–23^. We found additional mutations unique to the host specialists in the gene *rph,* encoding RNase PH that has not been linked to c-di-GMP signaling previously. Using both a fluorescence-based c-di-GMP sensor and liquid chromatography-mass spectrometry (LC-MS), we found a roughly twofold c-di-GMP increase in three wrinkly isolates, each with a single mutation in either *wspE*, *wspF*, or *rph*, when compared to the ancestor (Fig. 2b, line 152, Extended Data Fig. 7, Supplementary Table 10). This points to a loss-of-function mutation in *wspF* (which downregulates c-di-GMP) and alterations of active sites of WspE and Rph that all converge at upregulating c-di-GMP. We subsequently asked whether these specific mutations indeed cause improved host-association.

A functional genetic analysis of *wspE*, *wspF*, and *rph* demonstrated their direct involvement in host adaptation. For this analysis, we assessed the competitive fitness of mutants relative to the ancestor during host colonization. First, we re-assessed the three selected wrinkly mutants and found them to be significantly more competitive than the ancestor (Fig. 2c, left panel, line 152, Supplementary Table 11). Thereafter, we rescued these mutants with the corresponding ancestral alleles, which indeed abolished the mutants’ fitness increase (Fig. 2c, middle panel, line 152, Supplementary Table 11). Thirdly, an experimental introduction of each mutation into the ancestral background resulted in a significantly higher competitiveness, at least for the *wspF* and *rph* mutations (Fig. 2c, right panel, line 152, Supplementary Table 11). A similar fitness advantage was observed for the *wspE* and *wspF* mutants when either was subjected to quartet competition with the ancestor and the two other morphotypes (Extended Data Fig. 8, Supplementary Table 12). We conclude that changes in *wspE*, *wspF*, and *rph* that converge on increasing c-di-GMP levels enhance bacterial fitness in the host. As upregulation of this second messenger mediates a fundamental life history switch^13^, we next investigated whether it more generally mediates host-association across Pseudomonads.

### c-di-GMP generally promotes symbiosis

Genetic manipulation of *wspF* and a bioinformatic analysis of *Pseudomonas* genomes revealed a general involvement of *wsp* genes in host-association. For the former, we generated *wspF* deletion mutants for *P. lurida* strain MYb193 and the distantly related *P. alkylphenolia* MYb187 (both naturally associated with *C. elegans*), and further obtained mutant and wildtype *P. fluorescens* strain SBW25, a model for wrinkly formation^21^. We found that the mutants had significantly higher competitive fitness in the *C. elegans* host than their respective wildtypes (Fig. 3a, line 179, Supplementary Table 13). Furthermore, we correlated the presence of *wsp* and *rph* genes in 1359 whole *Pseudomonas* genomes from NCBI with the bacterial isolation source, a proxy for lifestyle (Extended Data Fig. 9, Supplementary Table 14). *Pseudomonas* isolates containing any of the genes or the complete *wsp* operon were significantly more often isolated from a host than isolates lacking these genes (Fig. 3b, line 179, Supplementary Table 15). We propose that the presence of these genes allows regulating c-di-GMP and thereby to adjust to a host-associated lifestyle.

## Discussion

Together, our study demonstrates that bacteria can improve their association with a host by shifting their life history from a motile to a sessile, persisting lifestyle. This lifestyle shift results from correlated changes in a suite of life history traits (Fig. 1f, line 102), which together represent a transition in life history strategy. One way to interpret this transition is as a shift along the r-K life history continuum, from an r-like strategy characterized by high reproductive rates to a K-like strategy characterized by persistence under high density conditions^24, 25^. To demonstrate whether such a transition would generally lead to increased host-association, we used an extension of a previously published mathematical model of microbial evolution toward host-association^26^. Exploration of a broad parameter space with this model confirmed that increased within-host persistence is often the optimal strategy for microbial adaptation to hosts (Extended Data Fig. 10, Supplementary Discussion), suggesting that the results from our study may be generally applicable.

In our experiments, the lifestyle shift from primarily free-living to host-association is mediated by the universal second messenger c-di-GMP. C-di-GMP is well known to regulate key physiological functions in bacteria, including the regulation of virulence in bacterial pathogens^22, 27^. Our work demonstrates that this regulatory system promotes the adaptation of Pseudomonads to diverse host systems, from plants to human, not only in pathogens, but extending to beneficial host-bacterial relationships. Given the importance of beneficial microorganisms in the functioning of their hosts, understanding the mechanisms that mediate non-pathogenic associations is crucial. Our study suggests that c-di-GMP plays an essential role in many such associations.

## Methods

### Host and bacterial strains

We performed evolution experiments with *Pseudomonas lurida* strain MYb11 (*Pl*_MYb11) and its natural host *Caenorhabditis elegans* strain MY316 (*Ce*_MY316)^5^. In preparation of all experiments, we thawed frozen worm stocks (−80°C) and raised worms on nematode growth medium agar (NGM^28^) seeded with *E. coli* OP50. A standard bleaching protocol was used to collect sterile and synchronized L1 larvae, which were then raised to L4 stage on *E. coli* OP50 (20°C), unless stated otherwise.

*P. lurida* strains MYb11 and MYb193, and *P. alkylphenolia* MYb187 were isolated from *Ce_*MY316^5^, *P. fluorescens* SBW25 from sugar beet leaves^9^. Bacteria were cultured on tryptic soy agar (TSA; 20°C, 48 hours) and tryptic soy broth (TSB; 28°C, 150 rpm, overnight) unless stated otherwise.

### Evolution experiment

Bacterial populations originating from a clone of *Pl*_MYb11 were serially passaged on NGM in presence of *Ce*_MY316 (host treatment, 6 replicates) or without worms (negative control, 6 replicates). For each replicate, a lawn of *Pl*_MYb11 was seeded onto NGM and cultured for 3.5 days. For each cycle of the host treatment, ten *C. elegans* L4 larvae were added per plate and incubated until the worms reached the F1 generation (3.5 days). In the negative controls, bacteria were maintained on NGM without worms. At the end of every cycle, bacteria were collected from either worms or plates in the host-associated and control treatments, respectively, 10% of the population (bottleneck) was transferred to the next cycle, and a sample frozen (−80°C). A similar number of colony forming units (CFU) was used to bottleneck the negative control. A total of 10 cycles were performed.

Frozen bacteria from cycle 10 were recovered and before further experiments were conducted these were subjected to one more cycle of the evolution experiment to minimize any potential selective effects of freeze/thawing. To focus on evolved differences between populations of the host treatment and the negative control, rather than physiological responses to recent host exposure, bacteria were grown on NGM for two days as a common garden treatment and then used in subsequent assays.

### Bacterial colonization of individual worms

Bacterial fitness during host-association was quantified as CFU per worm. In preparation, bacterial lawns (125µl, OD_600_=2) were seeded on NGM and five synchronized L4 *Ce*_MY316 added. After 3.5 days at 20°C, worms were collected with M9 buffer containing 0.025% Triton-100 and 25mM of the paralyzing antihelminthic tetramisole. The worms were washed in buffer using a custom-made filter tip washing system^29^ and collected in M9 with Triton-100. Worm-free supernatant was collected as a background sample. Following homogenization by bead beating, serial dilution and plating were used to quantify CFUs. CFU/worm was calculated as the difference in CFU between worm and supernatant samples, divided by the number of worms per population. For diversified populations, colony morphologies were scored as smooth, fuzzy or wrinkly.

### Worm population growth

Worm population growth resulting from 5 L4 larvae over 3.5 days was quantified as a proxy for host fitness. Bacteria and worms were prepared as for colonization assays and washed worms frozen in 48-well plates. Photographs of worms were automatically scored in ImageJ2^30^: worms were detected as particles, approximated by ellipses, and those fitting *C. elegans*-like dimensions (major axis: 0.18-1.3 mm, minor axis ≤0.1 mm^31^) were counted. Detection quality was validated by correlating automatic worm counts with counts of two independent experimenters (*r*(58) = 0.736, *p* = 2.106 x10^-11^).

### Early colonization, persistence, and release in worms

To quantify early colonization, persistence, and release from L4 stage worms, bacterial lawns were prepared from ancestral *Pl*_MYb11 and evolved populations (post common garden) or clonal morphotypes (overnight cultures). In early colonization assays, we quantified bacteria that entered L4 *Ce*_MY316 that were previously raised on non-colonizing *E. coli* O50. Colonization levels were then assayed as above resulting in CFU/worm as a measure of early colonization.

For persistence and release assays, worms were raised on the respective assay bacteria (from L1 until L4 stage), mimicking the development of worms in the F1 generation of the evolution experiment. Worms were then collected, washed using the filter tip washing system and samples divided into supernatant (supernatant 1) and worm sample (100µl each). Worms were then suspended in 200µl M9 and incubated for 1 h, after which 100 µl supernatant containing released bacteria (supernatant 2) was collected. The CFU released per worm was determined by the difference in CFU in supernatant 2 and supernatant 1. Alongside, we quantified CFUs maintained in worms of this sample as a measure of persistence.

### Bacterial growth, colony expansion and swarming

To measure bacterial growth, bacterial populations (common garden treatment or overnight cultures) were adjusted to OD_600_=0.1 and 50 µl spotted to NGM. After incubation (24h or 3 days at 20°C), lawns were scraped off, homogenized and CFUs determined by serial dilution.

Colony expansion and swarming were assayed on NGM containing 0.5% or 3.4% agar, respectively. In either case, 0.5µl of cell suspension (OD_600_ = 1) was spotted on surface-dried agar plates. Colony diameter was measured after 24h, 3 and 7 days.

### Biofilm formation

*In vitro* biofilm formation was assayed in microtiter plates based as described previously^32^. Notably, assays were performed in a randomized layout in Nunclon Delta surface-treated plates. Staining was performed after 48 hours of incubation (20°C, orbital shaking: 180 rpm). Absorption of dyed biofilm solutions was measured at 550 nm.

### Isolation of morphotypes

Representative colonies with visually distinct morphologies were isolated from evolved cycle 10 populations. The evolved populations were thawed, serially diluted, and plated (48h, 20°C). Unique morphotypes from all evolved populations were re-streaked and archived as frozen stocks (Supplementary Table 3). All morphotypes were thawed and re-streaked once, and showed stable colony morphology for 2 days of incubation.

### Fluorescent labeling of wrinkly morphotype MT12 and *in vivo* microscopy

The wrinkly morphotype MT12 was labeled with red fluorescent dTomato (dT) using Tn7 transposon-based chromosomal insertion as previously described^33, 34^. Insertion of the label did not affect the wrinkly morphology of the colonies.

Fluorescently labeled MT12 was used to localize colonization in Ce_MY316 using confocal laser scanning microscopy (ZEISS LSM 880). For this, synchronized L1 stage larvae were exposed to labeled bacteria for 72h (20°C), then collected using gravity washing and mounted for microscopy as described before^34^. Overviews of complete worms were created using a 25× LD LCI Plan-Apochromat multi-immersion objective (NA=0.8; with Immersol^TM^ W (2010)) and details imaged using a 40× C-Apochromat water immersion objective (NA= 1.2). Bacterial fluorescence and worm autofluorescence were sequentially excited (561 nm and 488 nm), and detected with an Airyscan detector (R-S sensitivity mode; longpass filter: ≥ 570 nm; bandpass filter: 495–550 nm). Data was processed with the automatic Airyscan processing function of ZEISS Efficient Navigation 2. For a list of the genetically modified bacteria used in this study, see Supplementary Table 16.

### Genome sequencing and analysis

Total DNA was isolated using a cetyl-trimethylammonium-bromid-based (CTAB) protocol^35^. For Illumina HiSeq400 (paired-end, 300bp) sequencing, libraries were prepared using the Nextera DNA Flex kit. Read quality was inspected (FastQC v0.11.8^36^) and reads trimmed (Trimmomatic v0.3.9^37^). Paired reads were aligned to the MYb11 reference genome (RefSeq: GCF_002966835.1; Bowtie2 v2.3.3^38^) and duplicate regions removed (Picardtools v2.22.2^39^). Variants were called (BCFtools v1.10.2^40^ and VarScan v2.3.9^41^) and then annotated (snpEff^42, 43^). We filtered for non-synonymous variants not present in the ancestral control in R^44, 45^. Gene ontology was inferred using Pseudomonas.com^46^.

### Quantification of relative c-di-GMP abundances using a biosensor

To quantify intracellular concentrations of c-di-GMP in ancestral *Pl*_MYb11 and evolved wrinkly isolates (MT12: *wspF*_EVO_, MT14: *wspE*_EVO_ and MT22: *rph*_EVO_), we used an established plasmid-based biosensor^47^. Bacterial strains carrying the plasmid were grown on gentamicin-selective plates (70 h, 20°C). For microscopy, single colonies were resuspended in 1X PBS, spotted on 2% agarose patches on microscopy slides and sealed. Bacterial fluorescence was visualize using confocal laser scanning (ZEISS LSM 700 with 40× Plan-Apochromat oil immersion objective with NA of 1.4; Immersol^TM^ 518 F). Fluorescence of the sensor and normalizer were sequentially excited (555 nm and 488 nm) and detected with a photomultiplier tube detector (≤ 630 nm) and a variable secondary dichroic transmitting light (≤ 550 nm).

Fluorescence intensity per cell was measured in Image J^30^: all cells and five background areas were identified as regions of interest, and area, integrated density and mean gray values measured. This data from the untransformed images was used to calculate the corrected total cell fluorescence^48^.To infer c-di-GMP concentration, we calculated the relative fluorescence intensity (RFI), so the ratio between TurboRFP and AmCyan fluorescence intensities, as previously described^47^ and compared average RFIs between ancestral *Pl_*MYb11 and wrinkly morphotypes. For the images used in Fig. 2, linear LUT was used at full range. Brightness and contrast were applied equally to all images.

### Quantification of c-di-GMP using PRM LC-MS/MS

To quantify intracellular c-di-GMP using liquid chromatography-mass spectrometry (LC-MS) in parallel reaction monitoring (PRM) mode, ancestral and evolved *Pl*_MYb11 (MT12, MT14 and MT22) were grown in LB medium to an OD_600_ of 1.8 and pelleted by centrifugation. After washing with salt-free LB medium, pelleted cells were snap-frozen and stored (−80°C).

Cell were mixed with 10 pmol of internal standard (cyclic-di-GMP-^13^C_20_,^15^N_10_, Toronto Research Chemicals, Toronto, Canada) in 60 μL water. Extraction of c-di-GMP was performed as described^49^ with the following modifications: extraction solution (240 μL of 1:1 acetonitrile (ACN)/methanol (MeOH)) was added, and samples were vigorously vortexed. Following incubation on ice (15 min) and centrifugation (20,800 x g, 4°C, 2 min), extract supernatant was collected, and solvent extraction repeated twice (200 μL of 2:2:1 ACN/MeOH/water). Pooled extracts were dried, resuspended in 50 μL of water, and centrifuged to remove insoluble compounds. Concentrations of solubilized protein precipitates were determined using the Pierce BCA Protein Assay Kit (Thermo Fisher Scientific, Bremen, Germany). For LC-MS/MS, 1 μL extract was injected onto an EASY-nLC 1000 UHPLC (Thermo Fisher Scientific) and separated on a 15 cm ReproSil-Pur C_18_-AQ nano LC column (0.1 mm i.d., 1.9 μm, 120 Å, Altmann Analytik, München, Germany) at 400 nL/min. Eluent A was 10 mM NH_4_OAc, 0.1% HAc, eluent B was 100% MeOH. Chromatographic conditions were 5% eluent B (5 min), followed by a linear gradient from 5% to 20% B (15 min) and an increase to 70% B (1 min), followed by 70% B (5 min) and 5% B (5 min); HCD of the m/z 691.1021 and m/z 721.0714 precursors was performed on a Q Exactive HF Orbitrap MS (Thermo Fisher Scientific). Peak areas for the qualifying^50^ product ions m/z 248.0778 (light) and m/z 263.0965 (heavy) determined in Skyline^51^ were used to calculate total c-di-GMP amounts, which were normalized to total protein amount as obtained by the BCA assay.

### Mutant generation

A two-step allelic replacement method based on previously described protocols ^21, 52^ was used to introduce the evolved mutant alleles into an ancestral background and also to revert mutations by introducing ancestral alleles in the mutant background. We applied the following *modifications:* ∼700 bp long PCR amplicons surrounding each mutation were cloned into pUISacB (Extended Data Fig. 6) allowing for sucrose selection. The constructs were transformed into competent *E. coli* cells and transferred to *Pseudomonas* isolates via conjugative mating with an *E. coli* helper strain containing pRK2013^53^. Primers (see Supplementary Table 17) were designed using NCBI’s BLAST tool^54^ and NCBI Primer-BLAST^55^, NEBuilder v2.3.0 (New England Biolabs) and Oligo Analyse Tool (Eurofins Genomics). BLASTn and alignments with Clustal Omega^56^ were performed using default settings.

### *In vivo* competition assays

Competition experiments were performed as described for the short-term persistence assays. Co-inoculated bacteria were OD-adjusted and mixed in equal volumes before seeding as lawns on NGM agar. A *Pl*_MYb11 labeled with dTomato^34^ was used, which is equivalent to the ancestral *Pl*_MYb11, as no differences were observed in short-term persistence (ANOVA, F value = 0.99, degrees of freedom = 1, P value = 0.35). CFU/worm was determined by subtracting CFU in supernatants from those in worm samples. A competitive index was calculated as the ratio of CFU/worm of evolved or constructed mutants over CFU/worm of the ancestor.

### Correlation of *wsp* and *rph* gene presence with isolate source across Pseudomonads

Whole genome sequences from NCBI were mined for c-di-GMP modulating genes (focus: *wsp* operon, *rph*) with bacterial lifestyle in members of the genus *Pseudomonas*. First, candidate genomes were obtained (NCBI Nucleotide’s command line search tool; size: 5-8 mio. bp). This retrieved 2279 sequences, for which sample information from NCBI’s Biosample database was collected. When available, host, host disease status, isolation source and sample type were used to manually classify genomes as originating from free-living or host-associated isolates with or without/unknown disease (Data S3). Next, we downloaded all available *Pseudomonas* reference sequences for *rph* and *wsp* genes from pseudomonas.com^46^. These were used to identify candidate sequences of *rph*, *wspA*, *wspB*, *wspC*, *wspD*, *wspE*, *wspF*, and *wspR*. These target gene candidates were found in the selected genomes using BLAST (R package ‘rBLAST’) and filtered based on sequence lengths and percent identities of the BLAST hits (Extended Data Fig. 9). Percent identity and sequence length were selected to maximize the chance that genes were correctly identified (red rectangles in Extended Data Fig. 9). If at least one candidate gene was identified during BLAST searches with the reference genes as query, this gene was considered present in the respective genome. We then used χ^2^ goodness of fit tests to infer whether isolates with and without the target genes differed in the relative proportions of host-associated lifestyles (Supplementary Table 15).

### Statistical analyses

Before data analysis, assumptions of parametric models (normality, homogeneity of variances) were checked by visual inspection (box-/qqplots) and with Shapiro-Wilk and Levene tests. When these were not met, non-parametric tests were applied. Box-plots show median (center line), upper/lower quartiles (box limits) and 1.5x interquartile ranges (whiskers).

To check whether evolved populations differed from the ancestor in CFU/worm, we compared the shift in the evolved phenotype (ratio of CFU/worm of evolved populations over ancestral *Pl*_MYb11) to the ancestor using one-sample t-tests (alpha=0.05, mu=1) with false discovery rate (fdr;^57^) correction for multiple testing. We applied this approach to analyze: bacterial colonization of individual worms, worm population growth, early colonization, persistence, and release, colony expansion and swarming. To infer overall phenotypic shifts according to evolutionary treatment, a principal component analysis (PCA) including the assayed phenotypes was performed. We performed permutational analysis of variances (PERMANOVA, 1000 permutations) followed by pairwise comparisons of groups (fdr-corrected) to test for differences in phenotype sets of ancestral and evolved groups and plotted confidence ellipses (one standard deviation). Packages used: ggbiplot^58^, missMDA^59^, vegan^60^ incl. pairwise.adonis^61^.

Differences in proportions of the different colony morphologies (wrinkly, smooth and fuzzy) within worms were identified using generalized linear models (GLM; quasinormal distribution) with Tukey post-hoc tests (using lme4^62^, lmtest^63^, and multcomp^64^).

Changes in morphotype proportions over time were tested for using beta-regressions (using gamlss^65^).

Differences between morphotype phenotypes were detected using ANOVA or GLMs and followed by Tukey or Dunnett post-hoc tests. To infer functional specializations across phenotypes, we used PCA and PERMANOVA.

Differences in c-di-GMP concentrations between evolved isolates were inferred using Welch-ANOVA with Games Howell post-hoc comparisons.

We tested for differences in CFU/worm between morphotypes or mutants using GLMs and Dunnett or Tukey post-hoc comparisons.

All analyses and plotting were performed in R^44, 45, 66, 67^.

### Mathematical model

We built a model to assess the selection gradient experienced by bacteria during the evolution experiment (Extended Data Fig. 10). We focus on the phase when bacteria are in contact with worms, and consider a homogeneous population. The dynamics of the number of bacteria living in (any) host-association *n(t)* can be described by the equation

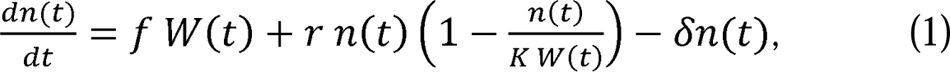

where *W(t)* denotes the biomass of worms on the plate at time *t*. We consider that growing to saturation on the plate are always in excess so that only the number of worms and the rate at which they feed on bacteria limit the immigration of free-living bacteria to the host: *f*. We assume logistic growth of the bacterial population within the worms, with maximal rate *r* and a carrying capacity proportional to the biomass of worms *W(t)* and the per unit of worm biomass carrying capacity *K*. Finally, a fraction of the host-associated bacterial population is removed from the host with rate δ, which encompasses bacterial death and expulsion to the environment. As in the evolution experiment, we assume that only host-associated bacteria are selected and continue to the next cycle, we ignore on-plate dynamics. We assume linear growth for the worm biomass, *W(t)=g t + W_0_*, encompassing both reproduction and development. We neglect the potential evolution of beneficial effects on worm growth and fix the parameters W_0_=10 and g=711 day^-1^ to experimentally observed values.

We study how the final number of host-associated bacteria, *n_f_*, is affected by changes in the parameters that describe the bacterial life cycle (*r,* δ*, f, K*). We define a range of biologically plausible values for each of these parameters (i.e. the trait space), that are informed by experimental data where possible:

– 10^-1^ day^-1^ < *r* < 10^1.25^ day^-1^, so between a small fraction and around twice the maximum on-plate growth rate (∼7 day^-1^).
– 10^-0.5^ day^-1^ *<* δ < 10^4^ day^-1^, as the typical time for a worm to lose 50% of its microbiome (in absence of feeding and replication) should range between seconds and days.
– 10^4^ < *K* < 10^6.25^, given the orders of magnitude from the maximal number of bacteria per worm measured experimentally (∼10^5^).
– 10^3^ day^-1^ < *f* < 10^7.5^ day^-1^, as the typical time for an empty worm to be colonized at 10% of its carrying capacity (*K*=10^5^); neglecting bacterial release and within-host replication) should be vary between seconds and days.

For each point of the trait space, we numerically solve equation (1) to compute the expected final number of bacteria at *t_f_*=3.5 days, *n_f_*=*n(t_f_)*. Finally, we assess the elasticity of *n_f_* along each direction of the trait space, which measures the expected relative change of *n_f_*with respect to a small relative change in one of the traits. We interpret the vector of the elasticities as the selection gradient on the phenotypic traits^68^ and use the dominant element of this vector to define an “optimal evolution strategy”^26^ for each point of the trait space.

## Data availability

Raw sequencing data is available in the NCBI Bioproject PRJNA862108. All other data is accessible via Dryad.

## Code availability

Custom code is available via Dryad, at https://github.com/nobeng and on reasonable request to the corresponding author.

## Acknowledgments

We thank E. Stukenbrock for LSM access; K. Guillemin, P. Rainey, H. Schweizer, H. Sondermann and F. Yildiz for providing bacterial strains; D. Rogers, J. Summers for guidance in allelic exchange; B. Pees for illustration support; S. Joel, J. Hofmann for lab support, the Kiel BiMo/LMB for access to their core facilities, the Schulenburg lab for project feedback, and B. Bohannan, R. Knight, P. Engel, and H. Sondermann for advice on the manuscript. Funding was provided by the Deutsche Forschungsgemeinschaft (DFG, German Research Foundation), Project-ID 261376515 – SFB 1182, Projects A4 and Z3 (NO, AC, FB, JL, A Tholey, A Traulsen, HS), the DFG Research Infrastructure NGS_CC project 407495230 (JF) as part of the Next Generation Sequencing Competence Network project 423957469, the International Max-Planck Research School for Evolutionary Biology (NO, AC), and the Max-Planck Society (Fellowship to HS).

## Author contributions

N.O.: conceptualization, methodology, investigation, analysis, writing, supervision. A.C.: conceptualization, methodology, investigation, analysis, writing. D.S.: analysis, writing. J.M.: investigation, writing. J.L.: investigation, writing. F.B.: conceptualization, modelling, analysis, writing. T.S.: investigation, writing. M.K.: investigation, writing. J.F.: investigation, writing. A.Th.: writing, supervision. A.Tr.: conceptualization, writing, supervision. H.S.: conceptualization, writing, supervision.

## Additional information

Supplementary Information is available for this paper. Correspondence and requests for materials should be addressed to Hinrich Schulenburg. Reprints and permissions information is available at www.nature.com/reprints.

## Extended data figures

**Extended Data Fig. 1.**
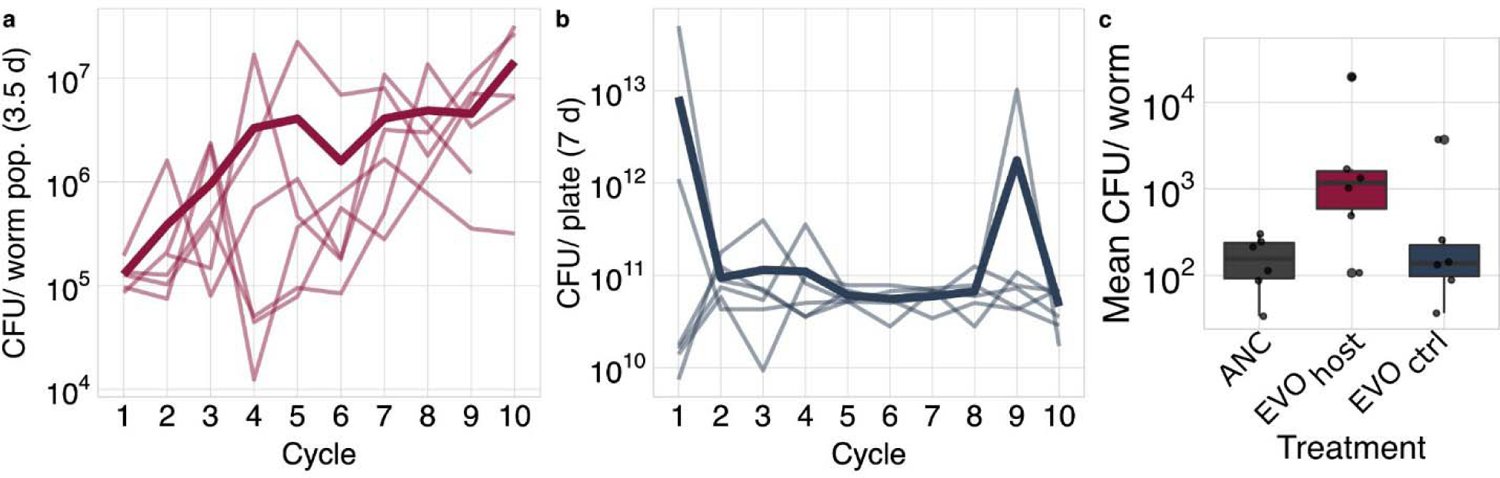
Bacterial fitness during the evolution experiment. a, Bacterial fitness in host across cycles of the evolution experiment measured as colony forming units (CFU) per worm population after 3.5 days of exposure to *Ce_*MY316. b, In the negative control, bacterial fitness was assessed on nematode growth agar in absence of the host. For each data point, bacteria were collected at the bottleneck time point of the noted cycle. Replicate populations (n=6) are shown as separate thin lines, with the mean shown as a thick line. c, Mean CFU per individual host in a worm population for the evolved bacterial populations of cycle 10. Five L4 *C. elegans* larvae proliferated on evolved or ancestral bacterial lawns for 3.5 days (reaching F1 generation) and CFUs were extracted from the whole worm population. CFUs per population were divided by the number of worms in the population. Overall results are shown as boxplots, with boxes indicating 25% above and below the median, which is given as the thick line within boxes; replicate populations are indicated as individual data points.

**Extended Data Fig. 2.**
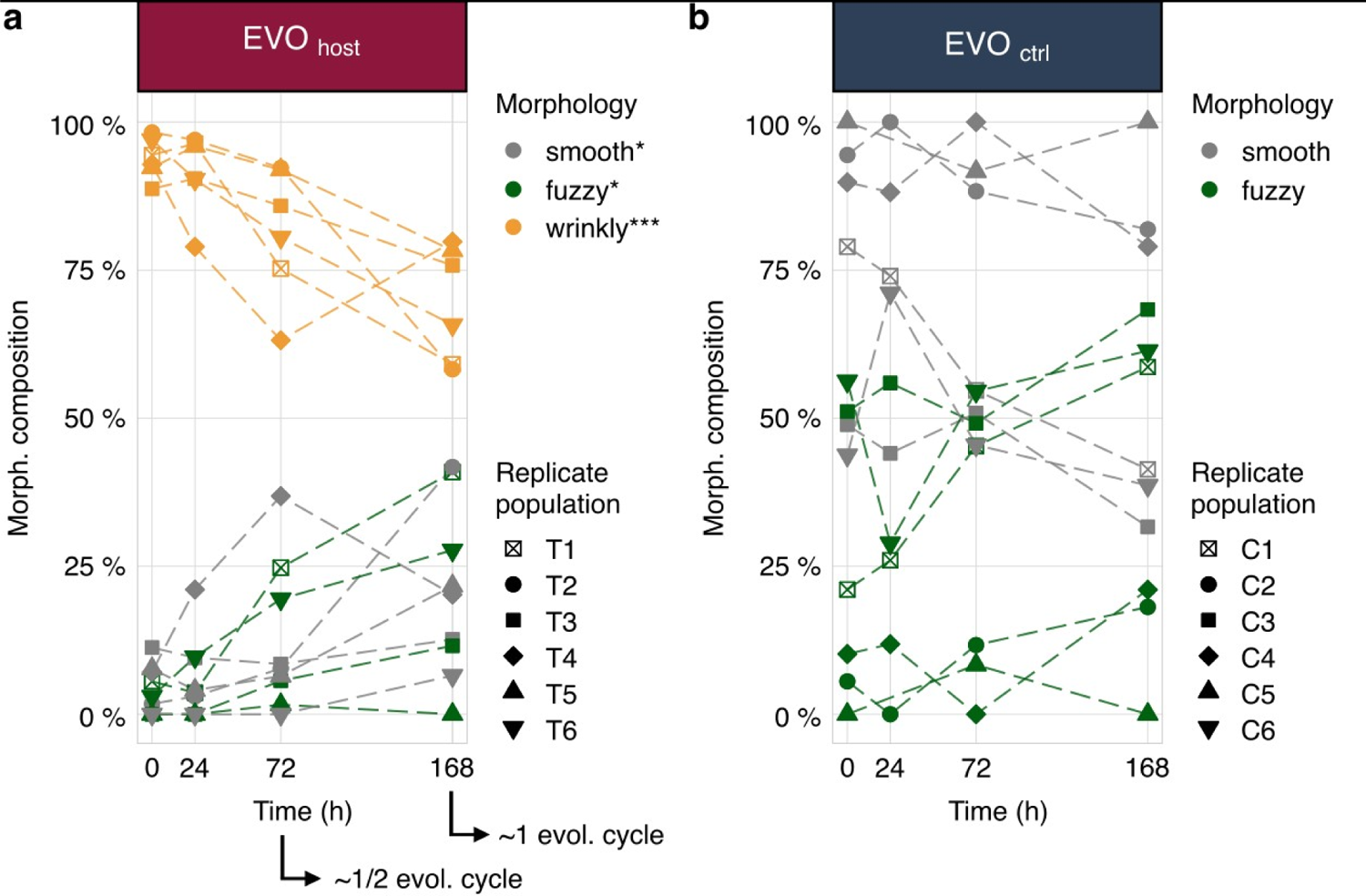
Dynamic changes in morphotype composition during the free-living phase of the host-associated life cycle for bacterial populations from the end of the evolution experiment. **a**, Results for the replicate populations from the host-associated evolution treatment. **b**, Results for the replicate populations from the control treatment. Proportions of the different colony morphotypes (see graphical legend) is shown across time of the host-associated life cycle. Time point 0 is at the end of the host-associated phase, when bacteria are transferred to the free-living phase, which itself lasts 168 hours. Beta-regressions were used to predict proportions and test for a change in proportions over time (fdr-corrected p-values: *** (p < 0.001), ** (p < 0.01), * (p < 0.05); see Supplementary Table 2).

**Extended Data Fig. 3.**
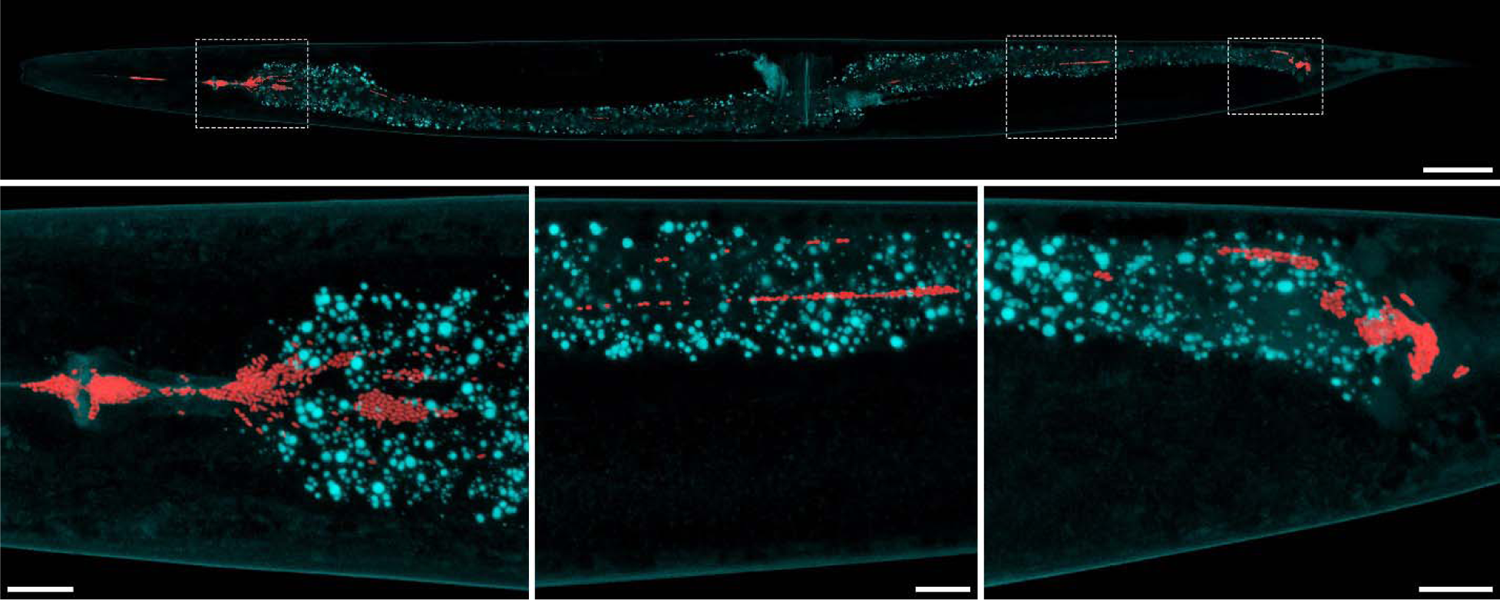
Colonization of the C. elegans intestine by wrinkly host specialists. Confocal laser scanning micrographs (maximum intensity projections) revealing intact bacterial cells (red) within the intestinal system of a young adult *Ce_*MY316 (cyan). The upper micrograph shows an overview of the complete worm, and the lower micrograph shows detailed views of the worm sections indicated by the dashed frames above. These include the posterior pharynx with the worm grinder and the first intestinal ring (left), a central intestinal (middle) and the anal region (right). The bottom left panel is identical to the micrograph shown in the main text (Fig. 1E, line 102). Scale bars = 50 µm (overview) and 10 µm (detailed views).

**Extended Data Fig. 4.**
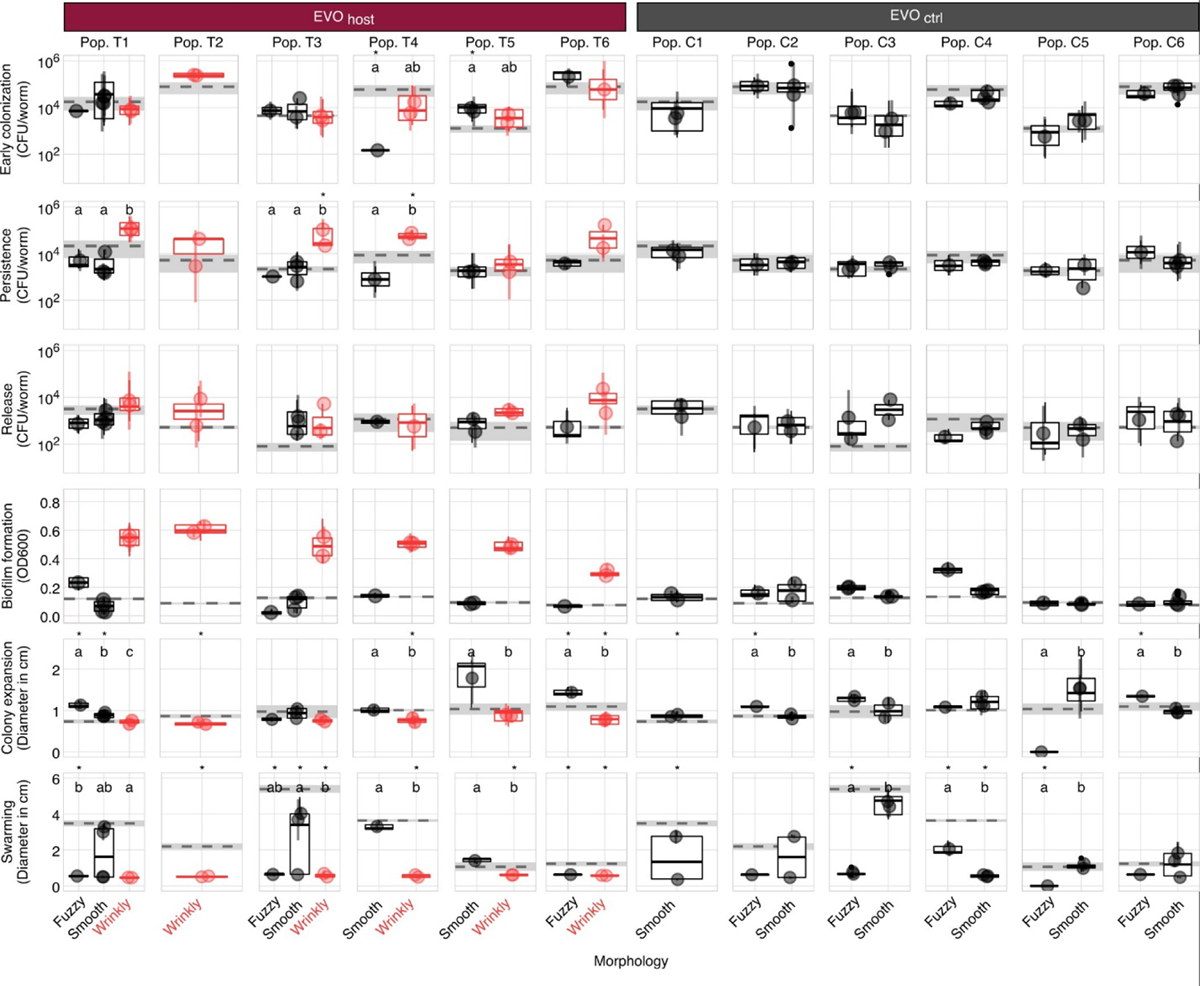
Wrinkly isolates from the end of the host-associated evolution treatment evolve a host-associated lifestyle. Phenotypes of morphotype clones isolated from independent host-evolved populations (left) and control populations (right), including smooth, fuzzy and wrinkly morphotypes, are shown. Results for each morphotype are summarized as boxplots. Dashed lines and grey shaded areas indicate the mean and standard error of ancestral traits, respectively. Differences between evolved morphologies and the ancestor were assessed with generalized linear models and Tukey post-hoc tests. Letters indicate statistical differences between morphologies, asterisks indicate deviation from the ancestor (see Supplementary Table 6).

**Extended Data Fig. 5.**
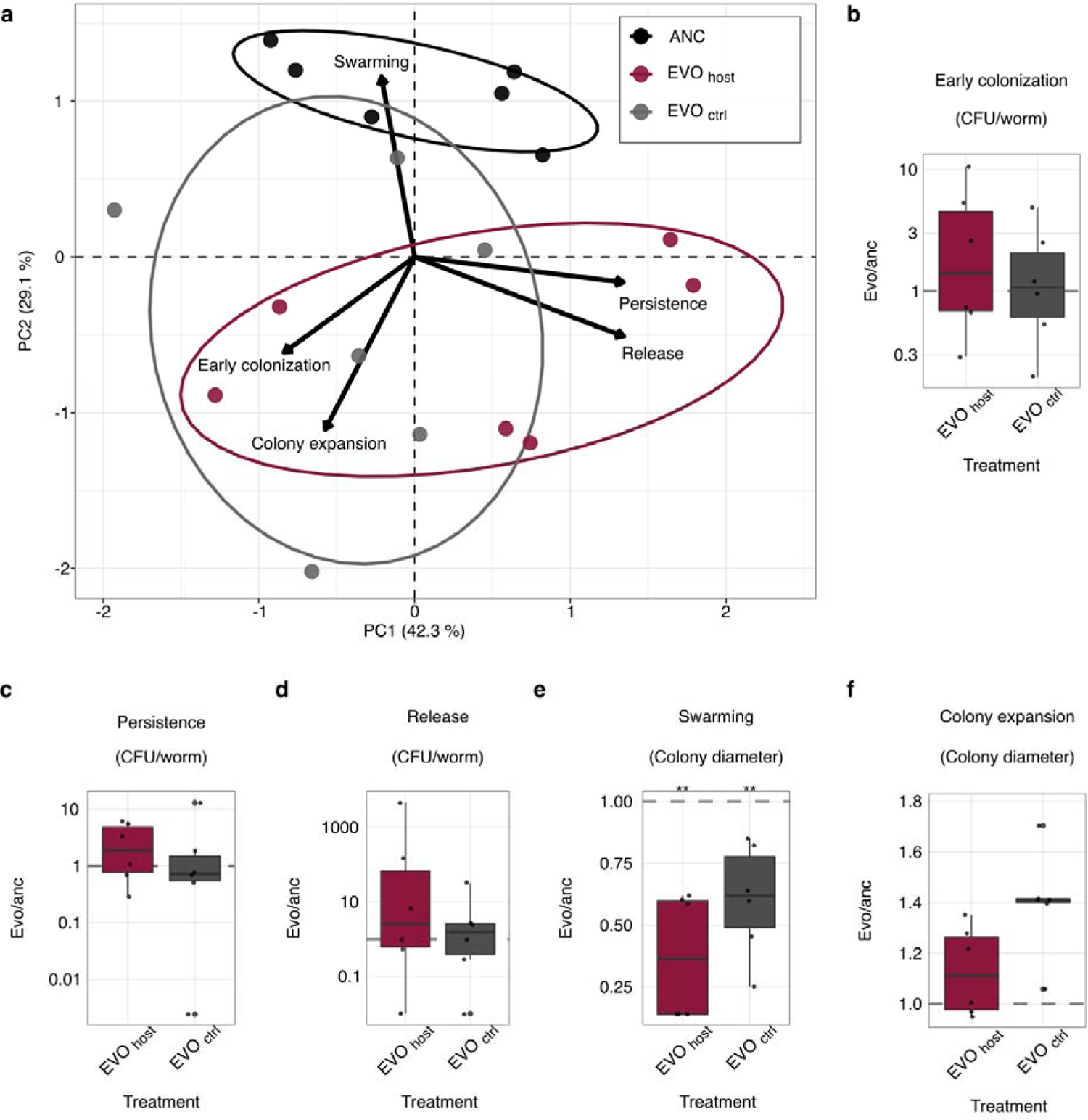
Evolution of a host-interaction life-style in the populations from the end of the host-associated evolution treatments. a, Principal component analysis on characteristic stages of host-association for ancestral, host evolved and control evolved bacterial populations. Individual data points refer to replicate populations colored according to evolution treatment (table S5). b-f, Shifts in phenotypes from the bacterial ancestor in the evolved populations for (b) early colonization, determined by CFU extracted from L4 larvae exposed to bacteria for 1.5 hour; (c) persistence in L4 larvae kept in M9 buffer for 1h (raised on bacteria); (d) CFU of bacteria released from L4 larvae into buffer within 1h (previously raised on bacteria from L1 to L4), (e) swarming distance on 0.5% agar within 24; and (f) colony expansion on 3.4% agar within 72h. All panels show ratios of evolved over ancestral populations for five replicates shown as individual data points. The dashed line indicates the mean values obtained for the ancestral population. The difference between evolved and ancestral phenotypes were assessed using t-tests (fdr-corrected; *** (p < 0.001), ** (p < 0.01), * (p < 0.05); Supplementary Table 8).

**Extended Data Fig. 6.**
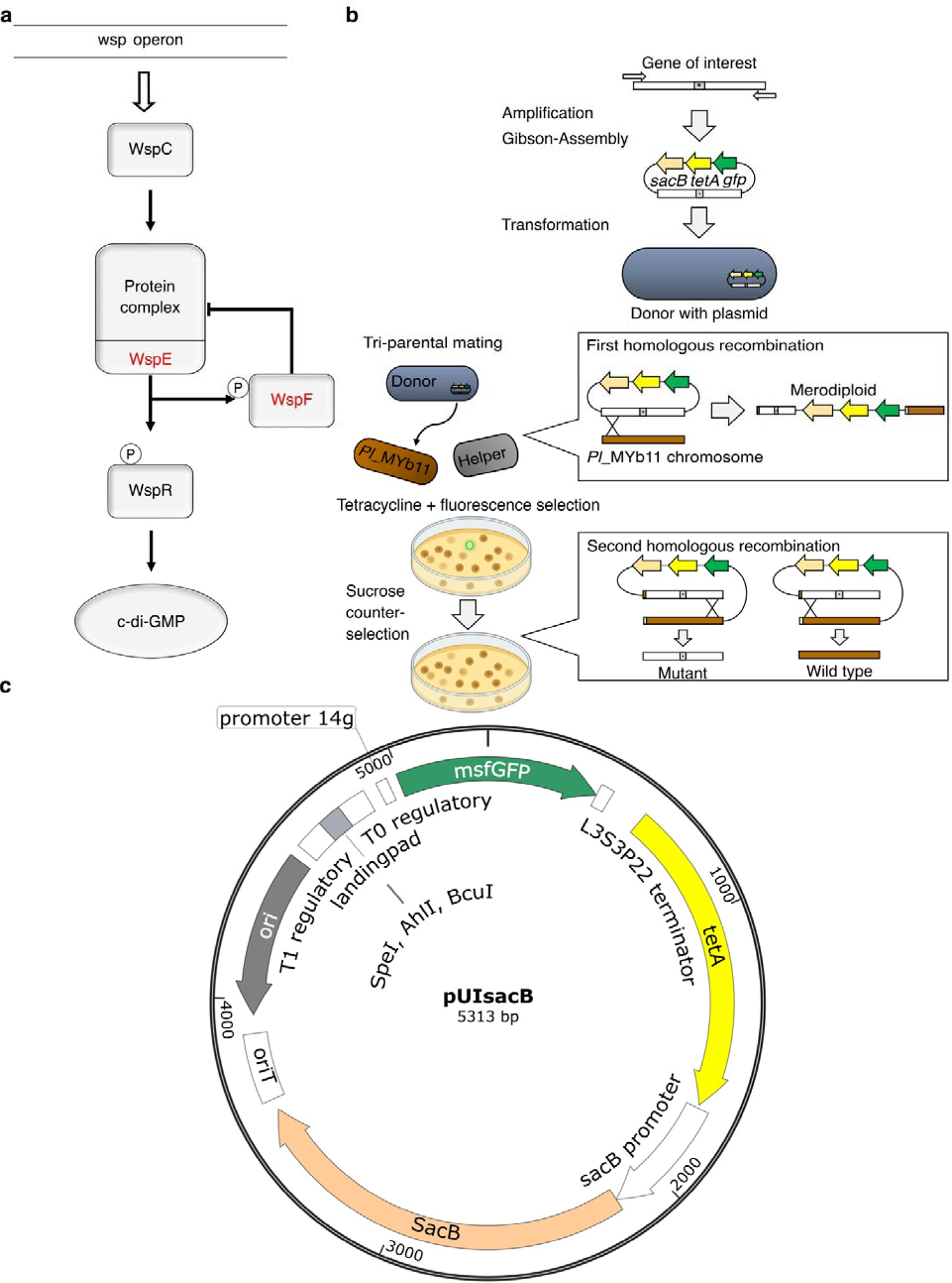
Allelic exchange strategy to generate Pseudomonas mutants in the wsp operon genes and in rph. **a**, Schematic representation of the wsp pathway of *Pseudomonas fluorescence* SBW25, as a model for c-di-GMP regulation in *Pl*_MYb11. The *wsp* operon encodes a two-component regulatory system that controls c-di-GMP production. Together, WspC, a methyltransferase, and WspF, a methylesterase, control the activity of the hybrid sensor histidine kinase WspE and subsequent c-di-GMP production by the di-guanylate cyclase response regulator WspR (after (18)). **b**, Workflow of the two-step allelic exchange strategy to introduce mutations into *Pl*_MYb11 (modified from (54), created with BioRender.com.). For conjugative pairing, **c**, pUIsacB (SnapGene software (from Insightful Science; available at snapgene.com)) containing an insert with ∼700 bp around each mutation, *tetA* (tetracycline resistance), *sacB* (sucrose sensitivity), and *gfp* was used. The first homologous recombination results in tetracycline-resistant fluorescent bacteria that are sensitive to sucrose. The second homologous recombination on sucrose results in loss of pUIsacB and insertion of the target mutation into the genome.

**Extended Data Fig. 7.**
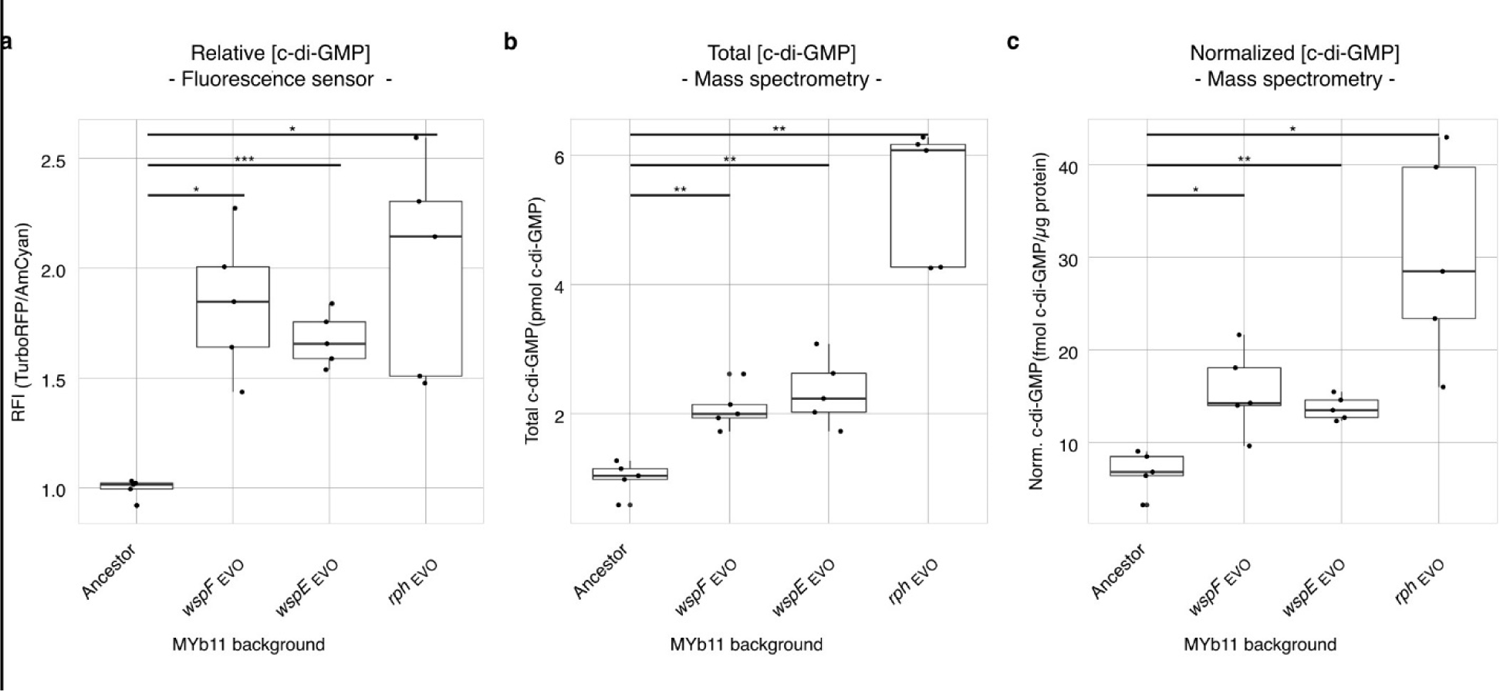
Increased intracellular c-di-GMP concentrations in wrinkly isolates. a, Amount of intracellular c-di-GMP measured with a fluorescence sensor. Raw fluorescence intensity (RFI) is the ratio of TurboRFP (c-di-GMP-dependent) and AmCyan (plasmid copy number-dependent) and, thus normalized for copy number of the sensor plasmid. b, Total c-di-GMP determined by isotope dilution PRM analysis. c, C-di-GMP determined by isotope dilution PRM analysis and normalized by total protein amount. In all cases, c-di-GMPs levels are studied for the ancestor and three wrinkly isolates from the end of the evolution experiment, each with a single mutation in either *wspF*, *wspE*, or *rph* (Supplementary Table 10).

**Extended Data Fig. 8.**
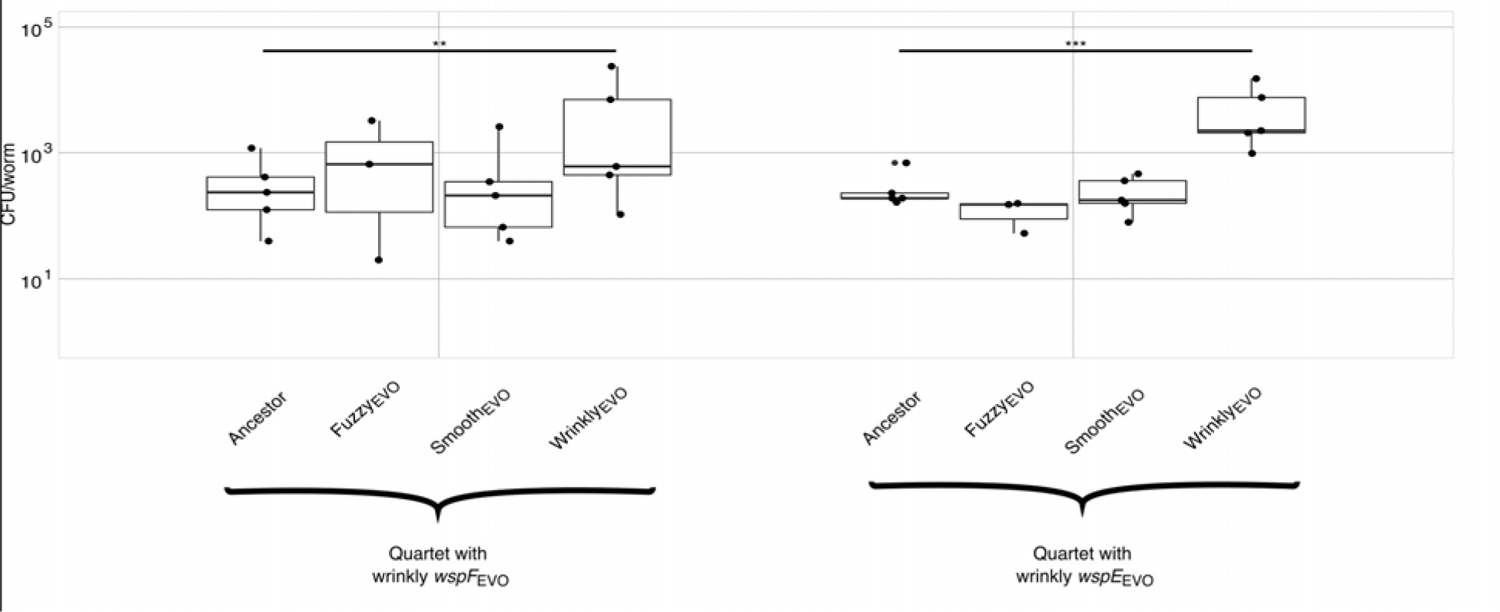
Increased competitive fitness of wrinkly isolates in bacterial mixtures of four strains (quartets). Three co-evolved morphotypes isolated from host-evolved replicate population T3 were paired with the ancestor. In one quartet the wrinkly *wspF* mutant (MT12) was present and in the other the wrinkly *wspE* mutant (MT14). Data points represent independent replicates, min. 3 replicates per treatment. Differences between morphotypes and ancestor were assessed using a linear model and subsequent Dunnett post-hoc comparisons; p-values: *** (p < 0.001), ** (p < 0.01), * (p < 0.05); see Supplementary Table 12.

**Extended Data Fig. 9.**
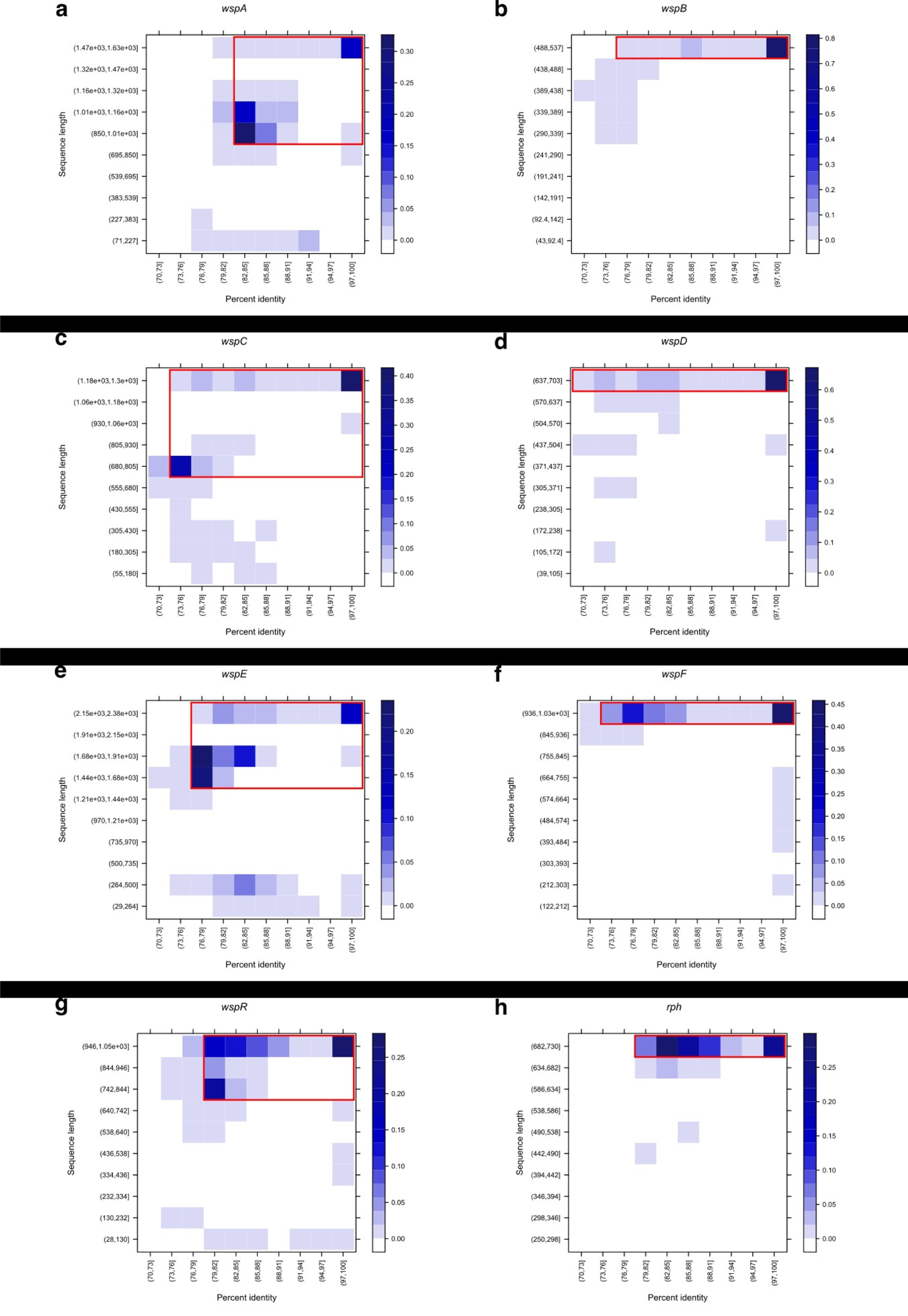
Filters for the identification of *rph* and *wsp* gene candidates. Distribution of sequence lengths and percent identities of the BLAST results for individual genes. The proportion of BLAST results belonging to particular sequence length and percent identity classes are shown as blue shades with varying intensity (cf. scale). Red rectangles show the areas for which the presence of the considered gene is assumed, and were set to include the largest BLAST hit values at both maximum sequence length and percent identities.

**Extended Data Fig. 10.**
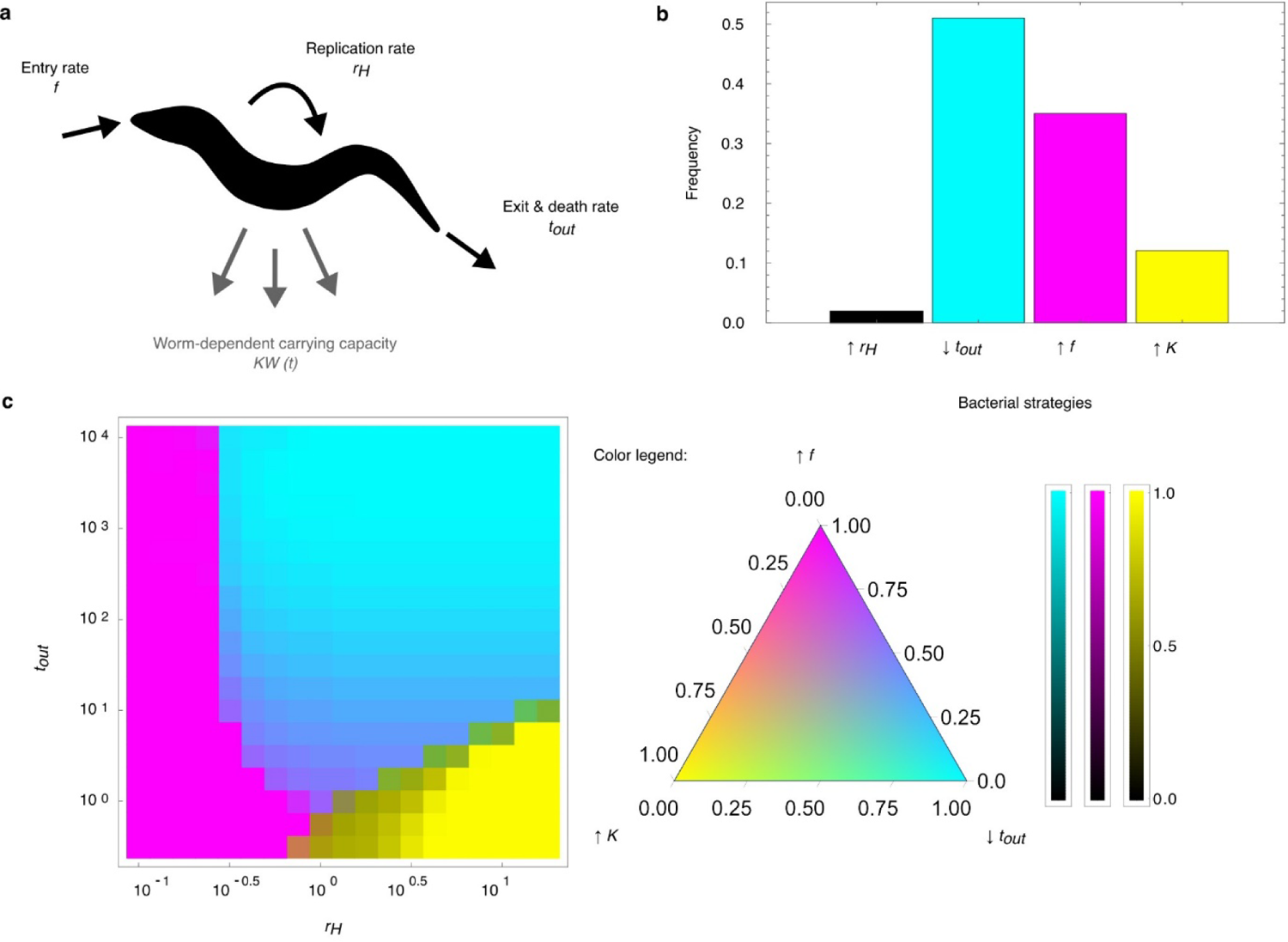
Model assessing the selection gradient on bacteria following a host-associated life cycle. **a**, Definition of the rates for the model of a microbial lineage being taken up by, replicating within and being expulsed from worms on a plate. **b**, Distribution of the optimal strategies across the whole traits space. **c**, Projection of the trait space over the axes (*r*, δ). For each point of the map, the color represents the proportions of times each of the 4 possible strategies are optimal, integrating over the values of *f* and *K*. The color scheme uses the CMYK color code: a purely cyan (respectively magenta, yellow) pixel indicates that the only optimal strategy for the considered values (*r,* δ) is ^—^ δ (respectively - *f*, - *K*). A color of a darker shade indicates that - *r* is also optimal in a small proportion at that point, as shown on the additional color scales for each edge.

